# DeepTrio: Variant Calling in Families Using Deep Learning

**DOI:** 10.1101/2021.04.05.438434

**Authors:** Lucas Brambrink, Alexey Kolesnikov, Sidharth Goel, Maria Nattestad, Taedong Yun, Gunjan Baid, Howard Yang, Cory Y McLean, Pi-Chuan Chang, Kishwar Shafin, Andrew Carroll

**Affiliations:** Google Inc, 1600 Amphitheatre Pkwy, Mountain View, California, USA

## Abstract

Every human inherits one copy of the genome from their mother and another from their father. Parental inheritance helps us understand the transmission of traits and genetic diseases, which often involve *de novo* variants and rare recessive alleles. Here we present DeepTrio, which learns to analyze child-mother-father trios from the joint sequence information, without explicit encoding of inheritance priors. DeepTrio learns how to weigh sequencing error, mapping error, and *de novo* rates and genome context directly from the sequence data. DeepTrio has higher accuracy on both Illumina and PacBio HiFi data when compared to DeepVariant. Improvements are especially pronounced at lower coverages (with 20x DeepTrio roughly equivalent to 30x DeepVariant). DeepTrio includes pre-trained models for Illumina WGS, Illumina exome, and PacBio HiFi.

## Introduction

Genomic sequencing can identify variants informative^1^ for diseases ^2^, traits ^3^, and ancestry ^4^. Sequencing is particularly informative in rare genetic disease^5^ caused by high-impact pathogenic variants ^6^. In a mother-father-child trio, each parent contributes half of their genome, with the addition of a small number of *de novo* variants ^7^. Sequencing for rare disease often includes the parents in order to use this information for accurate variant identification and interpretation^8^. Rare disease studies analyze multiple families with the same suspected disease to resolve undiagnosed cases^9^.

A number of methods can discover germline variants in an individual sample. Traditional methods model the evidence for a variant with known contributors to uncertainty, such as the rate of sequencing errors ^10^, the probability that a read is mapped incorrectly^11^, and the reliability of the sequence quality scores ^12^. Software tools which employ these statistical approaches include Freebayes ^13^, GATK ^14^, Octopus ^15^, 16GT ^16^, and Strelka2^17^.

Recent approaches have used deep learning, which learns representations directly from data ^18^. This allows a variant caller to capture aspects of the problem which are incompletely understood, or to rapidly adapt to a new sequencing technology by training on the latest data directly. Deep learning variant callers include DeepVariant ^19^, Clairvoyante ^20^, and Neusomatic ^21^. Deep learning was used in a majority of short-read and virtually all longread submissions to the PrecisionFDA Truth Challenge V2^22^.

Some variant callers can use information about a trio to jointly call variants. The approaches range from joint calling without consideration of family information (e.g. GATK GenotypeGVCF), those which model parental transmission probabilities (e.g, FamSeq^1^, GATK CalculateGenotypePosteriors), and those which use a deep learning approach for individual samples and postprocess with statistical methods (e.g. dv-trio ^23^).

Correctly incorporating parental information requires integrating the existing uncertainties in error sources across multiple samples. A deep learning approach can directly learn from trio data how corresponding evidence from parents may or may not support a call in the proband. It is also straight forward to adapt to different coverages, preparations, and technologies.

In this work, we build and assess **DeepTrio**, a deep learning-based variant caller for parent-child trios. We start from the code base of DeepVariant, a germline caller which won multiple awards in the PrecisionFDA Truth Challenge V2^22^, noted for high accuracy on genomes and exomes ^24^, and shown to increase detection rate of pathogenic germline variants ^25^.

We train DeepTrio to call variants in both parent and child samples, with one model for Illumina WGS and one for PacBio HiFi ^26^. To ensure accurate performance over a range of conditions, DeepTrio is trained with a diversity of preparations (PCR-free, PCR-positive), and across a range of child and parent coverages. DeepTrio can write output as individual VCFs ^27^, gVCFs, or as a merged family VCF. The gVCFs of multiple families can be combined with GLnexus ^28^, which has been optimized for combining DeepVariant gVCFs ^29^ to scalably create large joint callsets of trios and individual samples.

We show that DeepTrio has superior accuracy to both individual sample and trio-based samples, measured by concordance with the Genome in a Bottle truth set ^30,31^. We show that DeepTrio is able to accurately call *de novo* variants with high sensitivity despite their lack of support in the parents. Finally, we quantify the performance of DeepTrio across coverage, showing that DeepTrio allows high accuracy to be retained at lower coverage for both proband and parent samples.

## Results

### Modifying DeepVariant to call trios

DeepVariant calls variants in three steps: `make_examples, call_variants`, and `postprocess_variants`. In the `make_examples` step, a simple heuristic identifies positions which might differ from the reference. For Illumina sequencing, this requires a fraction of reads supporting an alternate allele at 0.12 for SNPs and 0.06 for Indels. For PacBio sequencing, the threshold is 0.12 for both SNPs and Indels.

The `call_variants` step represents BAM data as a multi-dimensional pileup of a 221- bp window in the genome around a candidate. As of DeepVariant v1.8, there are 9 possible input channels representing 1) the bases in the read, 2) their base quality, 3) the mapping quality of the read, 4) the strand mapped to, 5) whether the read supports a variant, 6) bases which differ from the reference. For Illumina data, we present 7) fragment length of the read, and in the case of PacBio data, 8,9) realignments of the reads to the alternate alleles. `postprocess_variants` converts the output probabilities of the neural network into a variant call and confidence, and resolves multi-allelic candidates into their most likely alleles.

To modify `make_examples` for trio calling, we perform candidate generation on each individual sample in the same manner. We also generate candidates from the union of reads from all samples with a reduced threshold for reads supporting the alternate allele, which allows discovery of alleles at a lower fraction but which are reinforced by appearing in multiple samples. When a candidate allele is identified in any single sample, it is also generated at the same position in the other samples. This allows us to generate an output variant probability for every candidate in every sample.

The `call_variants` stage uses a deep neural network to classify the probabilities for the genotype of each variant candidate. This process learns the important factors for classification directly from the input data. We generate tensor pileups where each sample has a fixed height with the child pileup in the middle, one parent’s pileup on top, and the other parent at the bottom. Using labels from Genome in a Bottle, we train two models: a child model and a parent model. To generate calls for each parent, two sets of examples are made, with a different parent as the top pileup.

The concept of Mendelian inheritance, or any explicit modeling of parent-child relationship is never provided to DeepTrio. Training simply creates these pileups and associated labels. The child model would learn that the reads in the middle pileup are most informative for calling, and the parent reads as supporting evidence. The parent model would learn that the topmost reads are most informative for the call, with child reads providing some supporting information, and the other parent marginal additional information.

The `postprocess_variants` stage is not altered. The final outputs are a VCF and a gVCF for each sample. Merging multiple gVCFs uses GLnexus ^28^ in a manner optimized for DeepVariant outputs ^29^. Figure 1 shows a representation of the calling process.

**Figure 1.**
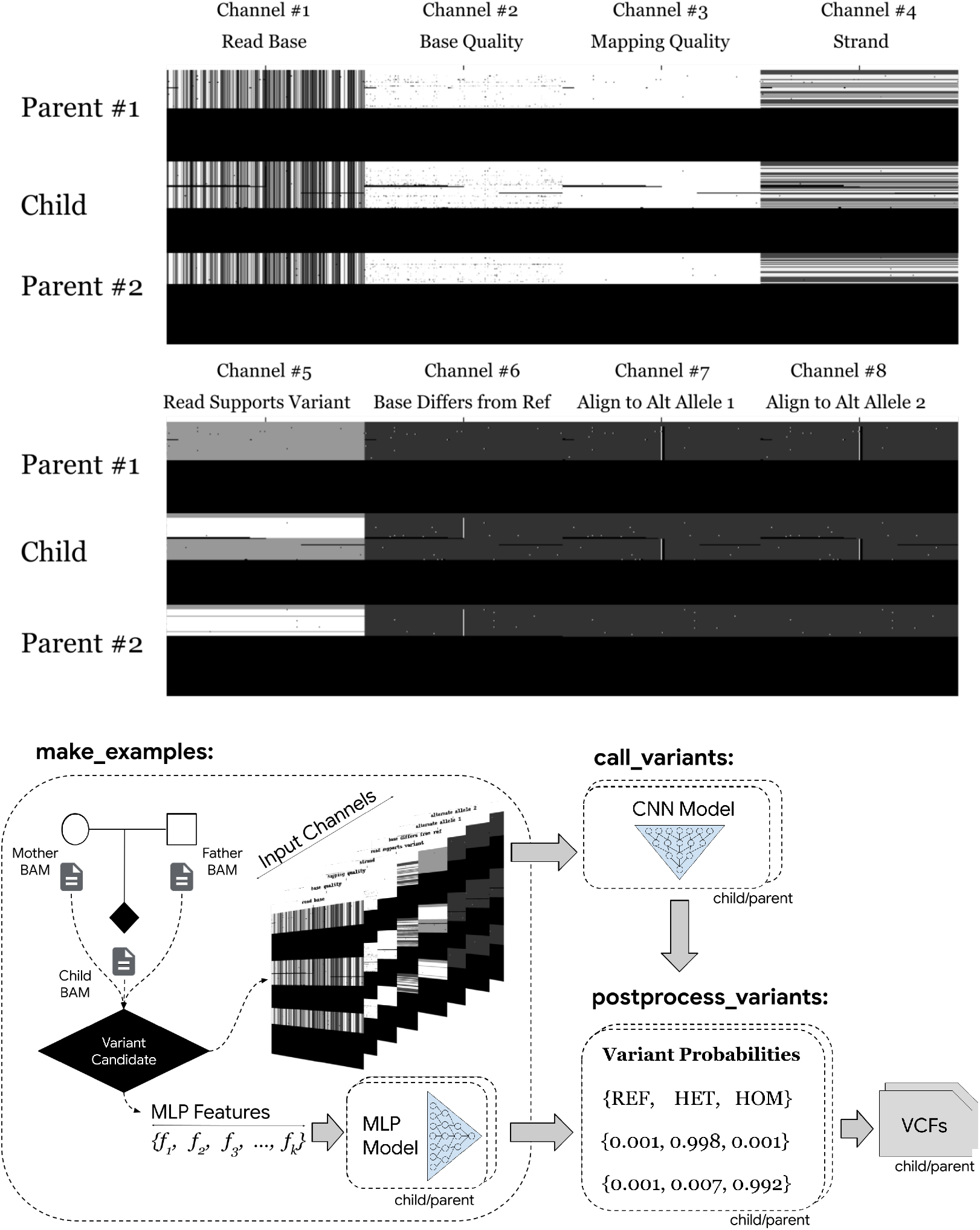
DeepTrio inputs channels and processing pipeline. DeepTrio represents data from the BAM file of a child and one or two parents as a pileup of a 221-bp long window (x-axis) with reads (y-axis). Each sample has a fixed height with parent and child reads in a different row. DeepTrio presents either 7 (Illumina) or 8 input channels (z-axis). Example shown is PacBio HiFi (top). One model is trained to call variants in the child and another in the parent. DeepTrio generates a distinct set of examples for each sample, alternating the position of the parents for the parent model, such that the parent sample being called remains in in the top row.

DeepVariant v1.8 introduces a multi-layer perceptron (MLP) classifier upstream of the primary variant caller. This light-weight classifier evaluates candidates via a 1D feature vector. Classifications that exceed a tuned confidence threshold bypass the deep neural network in `call_variants`, significantly reducing computational overhead. For DeepTrio, the 1D feature vector is adapted to encode variant- and allele information from all three samples. This performance optimization is particularly significant for DeepTrio due to the computational cost of tripling the height of the pileup images for trio analysis.

DeepTrio is trained on Genome in a Bottle samples^31^ with sequencing conducted on Novaseq, HiSeqX, HiSeq4000, and PacBio Sequel II instruments with both PCR-Free and PCR+ preparations. This data has been previously described and released^32^. DeepTrio is trained on examples from chromosomes 1-19. Chromosomes 21 and 22 are used as a tuning set to select the model checkpoint by determining when accuracy on the withheld set has peaked. Chromosome 20 is fully withheld as an independent evaluation set to assess accuracy.

### Assessing variant calling accuracy

To determine the improvements of trio calling, we compared DeepTrio to DeepVariant, GATK4 HaplotypeCaller (non-trio), GATK4 CalculateGenotypePosteriors (trio), dv-trio, and Octopus for the Genome in a Bottle (GIAB) Ashkenazi Jewish trio (HG002-HG003- HG004) datasets from the PrecisionFDA v2 Truth Challenge ^22^. Accuracy is determined by concordance with the GIAB v4.2.1 truth set^30,31,33^ using hap.py ^34^.

We observe that DeepTrio has higher accuracy for SNP and Indels than DeepVariant for both Illumina and PacBio HiFi data (Figure 2 top, Supplementary tables 1, 2), with a more pronounced effect at lower coverage depths (Supplementary tables 3, 4). For Oxford Nanopore long-reads, the trio-aware methods DeepTrio and Clair3-trio^35^ demonstrate high accuracy for SNPs (particularly below 20x), yet DeepVariant outperforms both methods on Indel calling across all coverages (Supplementary figure 8). This suggests that ONT’s systematic Indel errors are replicated across samples and thereby reinforce false evidence.

**Figure 2.**
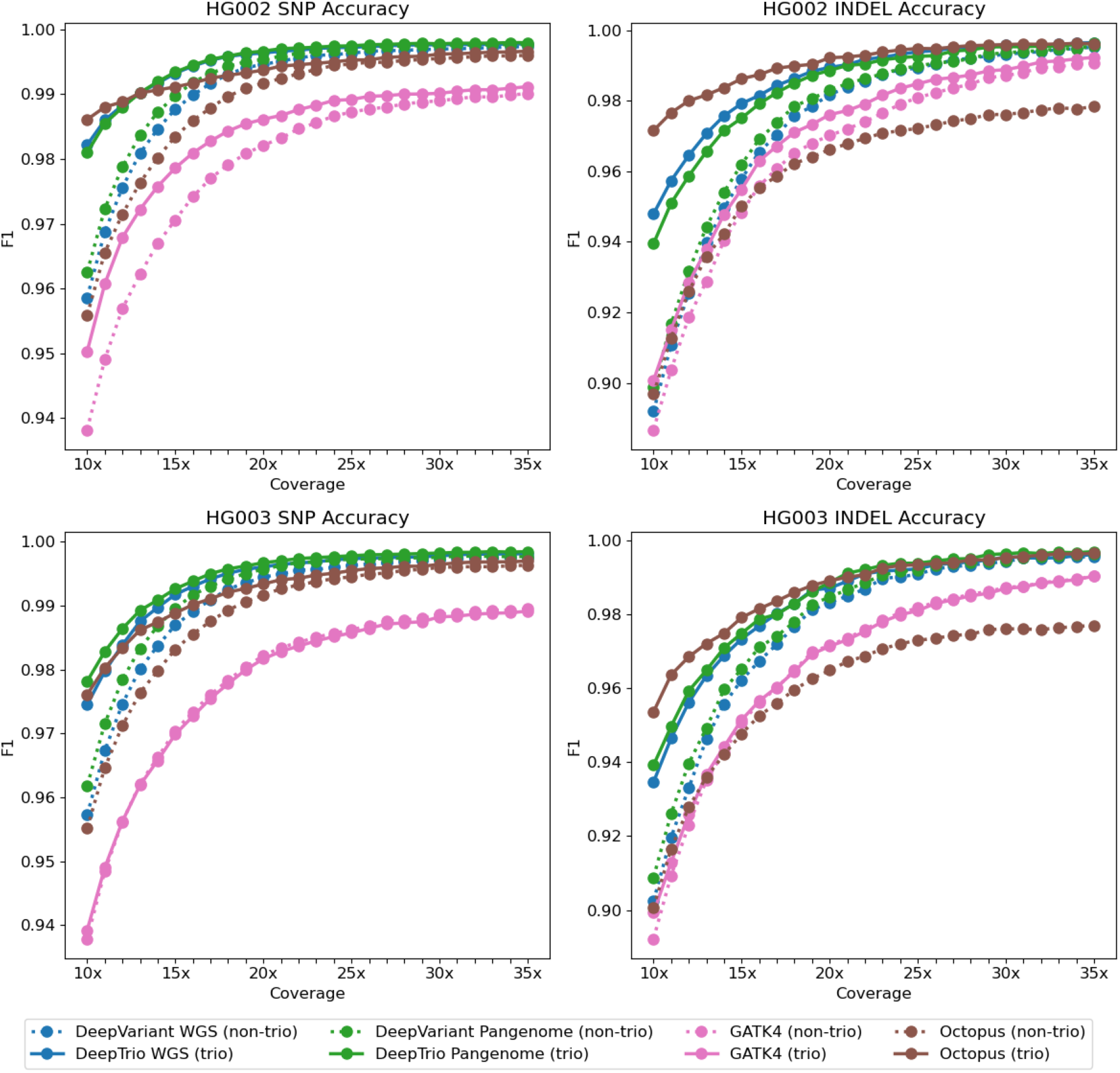
Variant calling accuracy across different pipelines and sequencing depths. Accuracy for DeepTrio and other pipelines over coverages, determined by concordance with the Genome in a Bottle v4.2 truth set for the child (top) and parent (bottom), with the same coverage on all three samples. All samples use the same Illumina data, except for the green lines (DeepVariant PacBio and DeepTrio PacBio). Trio methods are shown with a solid line and non-trio with a dotted line. F1 is determined from the total errors and correct calls for total Indels and SNPs for chromosome 20. The F1 for dv-trio was much lower and was excluded.

### Assessing variant calling accuracy in the parent samples

Accurate variant calling of parent samples is important in order to give context for proband variants, to accurately catalog incidental findings, and to correctly generate research cohorts. To assess the accuracy of parent calling, we compared DeepTrio’s parent model performance with DeepVariant and other trio and non-trio methods (Figure 2 bottom). DeepTrio outperforms other pipelines across a range of coverages, with a larger effect at lower coverages. DeepTrio’s advantage is less pronounced for the parent model as compared to its advantage on the child model. This finding is reasonable, since the genotype of the child is less informative regarding the genotype of the parent than the parent’s genotype is informative regarding the genotype of the child. The improvement in accuracy allows a lower coverage for the parent sample to be used while retaining the accuracy one would normally achieve with a non-trio pipeline.

### Precision and recall of *de novo* variants

The ability to identify *de novo* variants which may have a dominant inheritance pattern is of particular interest in identifying rare genetic disease. Because *de novo* variants violate the assumptions of Mendelian inheritance, it is reasonable to think that trio calling approaches may have a more pronounced effect on the sensitivity and specificity of *de novo* variants. When the analysis is constrained to cases where a child is called as 0/1 and each parent is confidently called as 0/0, we observe far more pronounced differences between trio and non-trio pipelines, even in cases like GATK4 where the overall accuracy as determined by Genome in a Bottle comparison is very similar.

The trio-aware pipelines (DeepTrio, dv-trio, GATK CalculateGenotypePosteriors), and Octopus have a greatly reduced false positive rate for *de novo* variants, but also demonstrate a reduced recall of true *de novo* variants, especially at lower coverages (Figure 3, Supplementary tables 1, 2, 3, 4). True *de novo* variants are defined by confidently genotyping the child as 0/1 and the parents as 0/0. DeepTrio is able to confidently identify *de novo* variants at lower read depths compared to other trio-aware pipelines evaluated (Figure 3 bottom).

**Figure 3.**
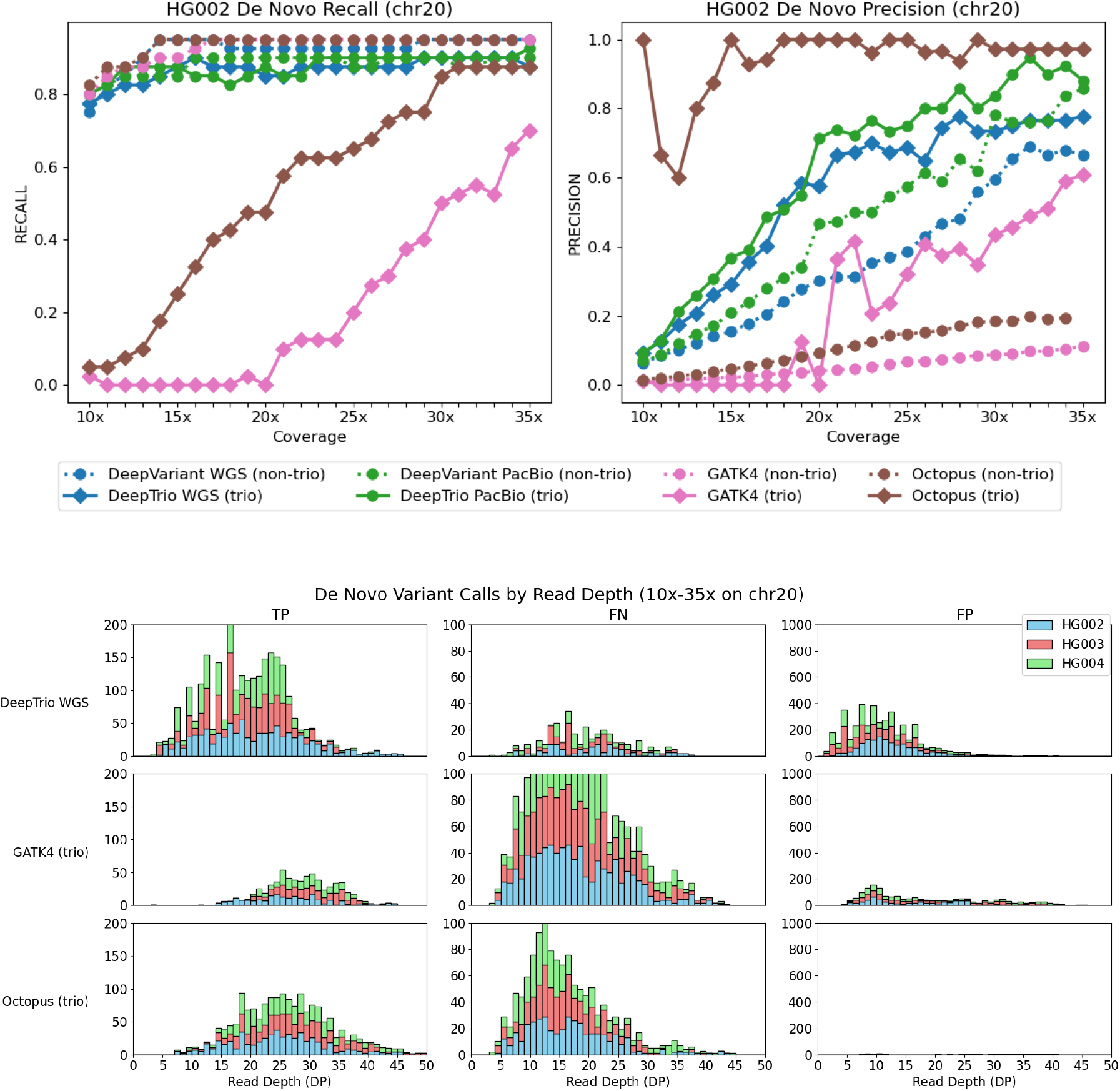
Performance of *de novo* variant calling across different pipelines and sequencing depths. (Top) Recall (left) and precision (right) for *de novo* variants (child *0/1*, parents *0/0*) on chromosome 20 (HG002) as a function of sequencing coverage. (Bottom) Breakdown of call accuracy (True Positives (TP), False Negatives (FN), and False Positives (FP)) by read depth for trio-aware pipelines (DeepTrio, GATK4, and Octopus) on Illumina WGS data. *De novo* sites are defined using Genome in a Bottle (GIAB) v4.2.1 benchmarks. DeepTrio demonstrates superior performance, particularly maintaining higher recall at lower read depths compared to GATK4 and Octopus.

### Assessing the effect of parental depth on variant calling accuracy in the proband

In trio sequencing for rare disease, there is often greater importance in sequencing a child proband. To manage sequencing costs, studies often take the approach of sequencing the parents at a lower coverage than the child^36^. In order to evaluate performance in these scenarios, we performed downsampling of the parent samples while keeping the child coverage at 35x. Variant calls were generated with the same tools used for the coverage titration of the full trio and evaluated using the same methods.

For the child sample, which contains the same reads for each titration point, we observe a slight accuracy improvement for DeepTrio in both Illumina and PacBio HiFi across the parent coverage ranges observed (10x-35x). For the parent samples, which do vary in coverage over the titration range, we observe a much greater advantage for DeepTrio and trio-aware methods compared to DeepVariant and other non-trio methods (Supplementary figure 4). For PacBio HiFi, the parent model has higher accuracy when the child is at 35x, as compared to when all three samples are coverage-titrated.

### Inspecting examples of variant calls improved by DeepTrio

In order to better understand cases where DeepTrio makes a correct call where other methods do not, we inspected IGV ^37^ images of positions which DeepTrio called correctly that were errors in DeepVariant. Since the F1 of SNP calling at 35x is already 0.9978, all inspected sites were difficult (low coverage, low mappability, presence of repeats and segmental duplications). Errors corrected by DeepTrio were often either through the ability to identify supporting evidence at low coverage and difficult to map regions (Supplementary figure 7), or the ability to better estimate the correct genotype in sites with substantial allelic bias (Supplementary figure 6).

### Computational efficiency of DeepTrio and other trio-calling pipelines

To assess the computational efficiency of DeepTrio compared to other methods of generating trio calls, we ran pipelines on the Illumina 35x WGS samples using the same hardware: a 16-CPU thread machine (n1-standard-16) available on Google Cloud Platform. This was chosen as representative of typical runs, though there are faster (and more expensive) or slower (but cheaper) methods to run these pipelines (an analysis of single-machine scaling can be found in the DeepVariant-GLnexus paper^38^).

Some components of DeepTrio are more computationally efficient when compared to DeepVariant, for example DeepTrio can reuse calculations in the example generation stage. Other components require more compute in DeepTrio: the size of the pileup images is larger, which corresponds to a larger neural network and more compute; however, the introduction of a light-weight secondary classifier in v1.8 eases this computational burden significantly. Also, more examples are identified across the samples and any example in one sample requires making a call in the other. We observe that DeepTrio requires more time to run compared to DeepVariant, but significantly less time than GATK4 or Octopus on the same hardware (Supplementary figure 8).

## Discussion

Here we discuss how DeepTrio’s performance characteristics relate to the motives for trio calling and the data which is generally available. In this work, we considered accuracy across all variants, and separately for *de novo* variants. In the context of rare disease, both of these formulations of accuracy are important. Some rare genetic diseases are caused by the combination of recessive variants from each parent. While others, especially those with a dominant inheritance pattern, will arise *de novo*.

We demonstrate that DeepTrio has strictly superior accuracy compared to DeepVariant and other trio and non-trio methods. When considering highest overall accuracy, DeepTrio would always be preferred. The case of *de novo* variants is more nuanced. DeepTrio has much higher recall when calling *de novo* variants but has slightly lower precision when compared to Octopus at 35x. This effect is especially pronounced below 30x coverages, where DeepTrio still manages to accurately identify *de novo* sites while other trio-aware pipelines such as GATK4 and Octopus do not. It should be noted that non-trio aware methods such as DeepVariant provide stillhigher recall still, albeit with a notable lack of precision. As a result, if investigators have a greater interest in the highest recall of *de novo* events, as opposed to overall accuracy, one option would be to run DeepVariant on each sample, and to re-run DeepTrio on the smaller number of regions where DeepVariant identified a *de novo* variant, in order to prioritize real variants first. In addition, we have previously shown that the output of the neural network is very well calibrated (Figure 2 of Poplin and et al. ^19^), and it may be possible to rank putative *de novo* events by the reported confidence of the call in order to tune DeepTrio more towards recall of *de novo*.

The accuracy analyses above are conducted at deep coverage (35x). Sequencing a trio of samples is more expensive than a single sample, and studies often compensate for this extra expense by sequencing at a lower coverage, especially for the parents. Reducing coverage is often considered when sequencing non-human disease samples, where the importance of accurately calling every variant is balanced against larger, more comprehensive studies. DeepTrio’s advantage over other methods is substantially greater at lower coverages. DeepTrio is about as accurate in 20x coverage as DeepVariant is at 28-30x coverage for both Illumina WGS and PacBio HiFi. DeepTrio is more accurate at 15x coverage than either GATK4 method at the highest coverage evaluated (35x). By maintaining high accuracy at reduced coverage, as well as by including training examples which reduce parent coverage while keeping child coverage, DeepTrio increases the flexibility of investigators to plan their trio sequencing to maximize cost-benefit.

As a deep learning method, DeepTrio does not explicitly encode the relationship between samples. DeepTrio’s ability to improve accuracy of calling, and to do so in a manner which is similar to human intuition regarding *de novo* variants, demonstrates an ability to capture rules which mirror general knowledge. This is similar to a recent demonstration which re-trained DeepVariant to use population allele frequencies ^39^. It is a strong indicator that deep-learning based variant callers can be further improved by finding ways to expose information which captures the underlying biology of samples and populations. Similarly, the framework of DeepTrio could, in theory, be further expanded to use sibling information, or to leverage more distant family relationships. Overall, the success of DeepTrio is a strong demonstration that thoughtfully identifying data which captures relevant biologi-cal or bioinformatics intuition, is a critical element to the development of strong machine learning methods in the genomics domain.

## Methods

### Generation of Sequencing Data

The generation of sequencing data for training is described in detail in Baid and et al. ^32^ and the WGS and PacBio evaluation data in Olson and et al. ^22^. In summary, all WGS and exome runs were conducted with 151-bp paired-end reads at 50x intended coverage from NovaSeq and HiSeqX platforms. For WGS, sequencing for both PCR-Free and PCR-Positive preparations. All sequencing was performed on HG001-HG007, NA12891, and NA12892.

For PacBio HiFi data, we requested 3 SMRT Cells 8M for each sample of HG003, HG004, HG006, and HG007. Libraries were prepared targeting a 15kb insert size and sequenced on Sequel II System with Chemistry 2.0. This was supplemented by data from Human Pangenome Reference Consortium (https://github.com/human-pangenomics/HG002_Data_Freeze_v1.0).

### Training models

Training DeepTrio requires a set of BAM or CRAM files and a set of truth labels. Examples are generated in the same manner used for calling and are annotated with the truth labels. A training tutorial for DeepVariant from input data is available at: (https://github.com/google/deepvariant/blob/r1.9/docs/deepvariant-training-case-study.md).

Using the trio data sets described in the prior section, DeepTrio was trained across a range of coverages achieved by random downsampling of 50x BAM files at fractions of 0.7 (35x), 0.5 (25x), and 0.3 (15x). This random downsampling helps DeepTrio to generalize across coverages, and we observe that having more difficult examples results in overall better models. Examples are generated for trios where each sample is downsampled at the same fraction, and those where only the parents are further downsampled while the child is kept at a higher coverage. Training occurs over the entire set of WGS samples to generate the WGS model.

Finally, to enhance the sensitivity of *de novo* detection, we implemented a targeted fine-tuning strategy by assigning a 50:1 class weight to candidate *de novo* sites. By disproportionately penalizing the misclassification of these rare events, we shift the model’s decision boundary to prioritize the recovery of *de novo* signals to remain sensitive to subtle allelic supports in the progeny that are absent in the parental pileups.

### Mapping and Variant Calling

Samples were mapped to GRCh3840 with BWA MEM ^40^ in an ALT-aware manner and deduplicated with Picard MarkDuplicates ^41^.

Variant calling was performed using DeepVariant v1.9.0, DeepTrio v1.9.0, GATK v4.2.6.0, Octopus v0.7.4, and dv-trio (DV+GATK+FamSeq). Timing estimates for DeepVariant used DeepVariant v1.1, using the OpenVINO acceleration by Intel, a recent contribution which speeds execution.

For DeepVariant, calling was performed following DeepVariant’s best practices in multi-sample calling (https://github.com/google/deepvariant/blob/r1.9/docs/trio-merge-case-study.md).

For GATK, non-trio aware calling was performed by HaplotypeCaller followed by GenotypeGVCFs. For GATK trio-aware calling, this VCF was further refined by CalculateGenotypePosteriors.

For dv-trio, single sample calling was performed as described in DeepVariant’s best practices in multi-sample calling, combined with GATK CombineGVCFs and then processed by FamSeq as outlined by the dv-trio methods. Since FamSeq only refines the call of SNPs, Indel calls were taken from the GATK VCF without modification.

For Octopus, single sample variant calling for single samples was performed using the v0.7.4 release, with the matched v0.7.4 germline forest model. The gVCFs were combined with GATK CombineGVCFs. Octopus trio calling was also done using the v0.7.4 release with the v0.7.4 germline forest model.

For all *de novo* analyses, only PASS entries were used for calculations across all callers. No call (./.) positions were excluded.

### Assessing Accuracy

Call sets were assessed using v0.3.9 of the haplotype comparison tool, hap.py ^34^. The v4.2.1 truth sets from GIAB^30,34^ were used to benchmark HG002-4 samples mapped to GRCh38. *De novo* sites were identified as heterozygous variants in HG002 absent in both parent truth sets within overlapping confident regions.

## Code and Data Availability

All DeepTrio code is available under a BSD-3 license at: https://github.com/google/deepvariant/tree/r1.9/deeptrio

All evaluation data are derived the Illumina and PacBio FASTQ files available from the PrecisionFDA v2 Truth Challenge22 at: https://precision.fda.gov/challenges/10

Training datasets are described in “An Extensive Sequence Dataset of Gold-Standard Samples for Benchmarking and Development”33 and download links for all sequence data are available in the supplement of that paper: https://www.biorxiv.org/content/10.1101/2020.12.11.422022v1.supplementary-material

## Acknowledgements

We thank Daniel Cook from Google for many helpful suggestions on the manuscript text and content. We thank Aaron Wenger and Billy Rowell from Pacific Biosciences for several helpful comments, especially suggestions to evaluate DeepTrio performance on chromosomeX.

## Author contributions

LB, AC, PC, and AK conceived and designed the study. LB, AK, SG, MN, GB, and PC wrote DeepTrio code. TY and CYM built and analyzed variant merging methods. LB, AK, PC, and AC analyzed results. AC and HY procured sequence data. AC and LB wrote the manuscript.

## Supplementary Material

**Table 1.** HG002 SNP accuracy at 35x Coverage (all trio members).

| Tool | Data Type | F1 | Recall | Prec | FN | FP | FP.gt | TP de novo | FP de novo |
| --- | --- | --- | --- | --- | --- | --- | --- | --- | --- |
| DeepVariant | Illumina | 0.9974 | 0.9954 | 0.9993 | 328 | 49 | 12 | 34/34 | 17 |
| DeepTrio (trio) | Illumina | 0.9978 | 0.9962 | 0.9993 | 270 | 48 | 8 | 33/34 | 10 |
| Octopus | Illumina | 0.9960 | 0.9938 | 0.9982 | 444 | 124 | 15 | 34/34 | 103 |
| Octopus (trio) | Illumina | 0.9966 | 0.9946 | 0.9986 | 382 | 97 | 16 | 32/34 | 1 |
| GATK4 | Illumina | 0.9901 | 0.9942 | 0.9861 | 413 | 1002 | 59 | 34/34 | 270 |
| GATK4 (trio) | Illumina | 0.9912 | 0.9943 | 0.9881 | 407 | 858 | 56 | 27/34 | 16 |
| dv-trio (trio) | Illumina | 0.9200 | 0.8774 | 0.9670 | 8745 | 2136 | 4 | 7/34 | 5531 |
| DeepVariant-PB | PacBio | 0.9996 | 0.9994 | 0.9997 | 43 | 20 | 14 | 34/34 | 1 |
| DeepTrio-PB (trio) | PacBio | 0.9997 | 0.9997 | 0.9998 | 24 | 17 | 12 | 34/34 | 0 |

**Table 2.** HG002 Indel accuracy at 35x Coverage (all trio members).

| Tool | Data Type | F1 | Recall | Prec | FN | FP | FP.gt | TP de novo | FP de novo |
| --- | --- | --- | --- | --- | --- | --- | --- | --- | --- |
| DeepVariant | Illumina | 0.9953 | 0.9931 | 0.9975 | 78 | 29 | 23 | 4/6 | 2 |
| DeepTrio (trio) | Illumina | 0.9965 | 0.9948 | 0.9981 | 58 | 22 | 14 | 2/6 | 0 |
| Octopus | Illumina | 0.9783 | 0.9731 | 0.9836 | 303 | 192 | 153 | 4/6 | 52 |
| Octopus (trio) | Illumina | 0.9962 | 0.9948 | 0.9976 | 59 | 28 | 7 | 3/6 | 0 |
| GATK4 | Illumina | 0.9907 | 0.9914 | 0.9900 | 97 | 117 | 43 | 4/6 | 34 |
| GATK4 (trio) | Illumina | 0.9923 | 0.9920 | 0.9926 | 90 | 87 | 39 | 1/6 | 2 |
| dv-trio (trio) | Illumina | - | - | - | - | - | - | - | - |
| DeepVariant-PB | PacBio | 0.9924 | 0.9926 | 0.9921 | 83 | 92 | 49 | 2/6 | 5 |
| DeepTrio-PB (trio) | PacBio | 0.9946 | 0.9958 | 0.9935 | 47 | 77 | 30 | 3/6 | 5 |

**Table 3.** HG002 SNP accuracy at 20x Coverage (all trio members).

| Tool | Data Type | F1 | Recall | Prec | FN | FP | FP.gt | TP de novo | FP de novo |
| --- | --- | --- | --- | --- | --- | --- | --- | --- | --- |
| DeepVariant | Illumina | 0.9946 | 0.9919 | 0.9973 | 576 | 193 | 94 | 34/34 | 68 |
| DeepTrio (trio) | Illumina | 0.9964 | 0.9946 | 0.9982 | 387 | 130 | 52 | 33/34 | 20 |
| Octopus | Illumina | 0.9917 | 0.9866 | 0.9970 | 957 | 214 | 59 | - | - |
| Octopus (trio) | Illumina | 0.9937 | 0.9908 | 0.9967 | 659 | 234 | 26 | 0/34 | 0 |
| GATK4 | Illumina | 0.9821 | 0.9889 | 0.9755 | 792 | 1775 | 237 | 34/34 | 697 |
| GATK4 (trio) | Illumina | 0.9884 | 0.9896 | 0.9872 | 740 | 914 | 194 | 0/34 | 0 |
| dv-trio (trio) | Illumina | 0.9946 | 0.9919 | 0.9973 | 576 | 193 | 94 | 0/34 | 0 |
| DeepVariant-PB | PacBio | 0.9986 | 0.9977 | 0.9995 | 165 | 39 | 26 | 34/34 | 18 |
| DeepTrio-PB (trio) | PacBio | 0.9994 | 0.9993 | 0.9996 | 53 | 29 | 23 | 33/34 | 7 |

**Table 4.** HG002 Indel accuracy at 20x Coverage (all trio members).

| Tool | Data Type | F1 | Recall | Prec | FN | FP | FP.gt | TP de novo | FP de novo |
| --- | --- | --- | --- | --- | --- | --- | --- | --- | --- |
| DeepVariant | Illumina | 0.9818 | 0.9749 | 0.9888 | 283 | 129 | 106 | 3/6 | 18 |
| DeepTrio (trio) | Illumina | 0.9895 | 0.9851 | 0.9939 | 168 | 71 | 56 | 1/6 | 5 |
| Octopus | Illumina | 0.9662 | 0.9560 | 0.9766 | 495 | 270 | 221 | - | - |
| Octopus (trio) | Illumina | 0.9922 | 0.9894 | 0.9950 | 119 | 59 | 20 | 0/6 | 0 |
| GATK4 | Illumina | 0.9702 | 0.9708 | 0.9695 | 329 | 356 | 153 | 4/6 | 143 |
| GATK4 (trio) | Illumina | 0.9775 | 0.9733 | 0.9818 | 300 | 211 | 143 | 0/6 | 0 |
| dv-trio (trio) | Illumina | - | - | - | - | - | - | - | - |
| DeepVariant-PB | PacBio | 0.9744 | 0.9730 | 0.9759 | 304 | 280 | 138 | 2/6 | 23 |
| DeepTrio-PB (trio) | PacBio | 0.9886 | 0.9892 | 0.9880 | 121 | 141 | 68 | 2/6 | 7 |

**Figure 4.**
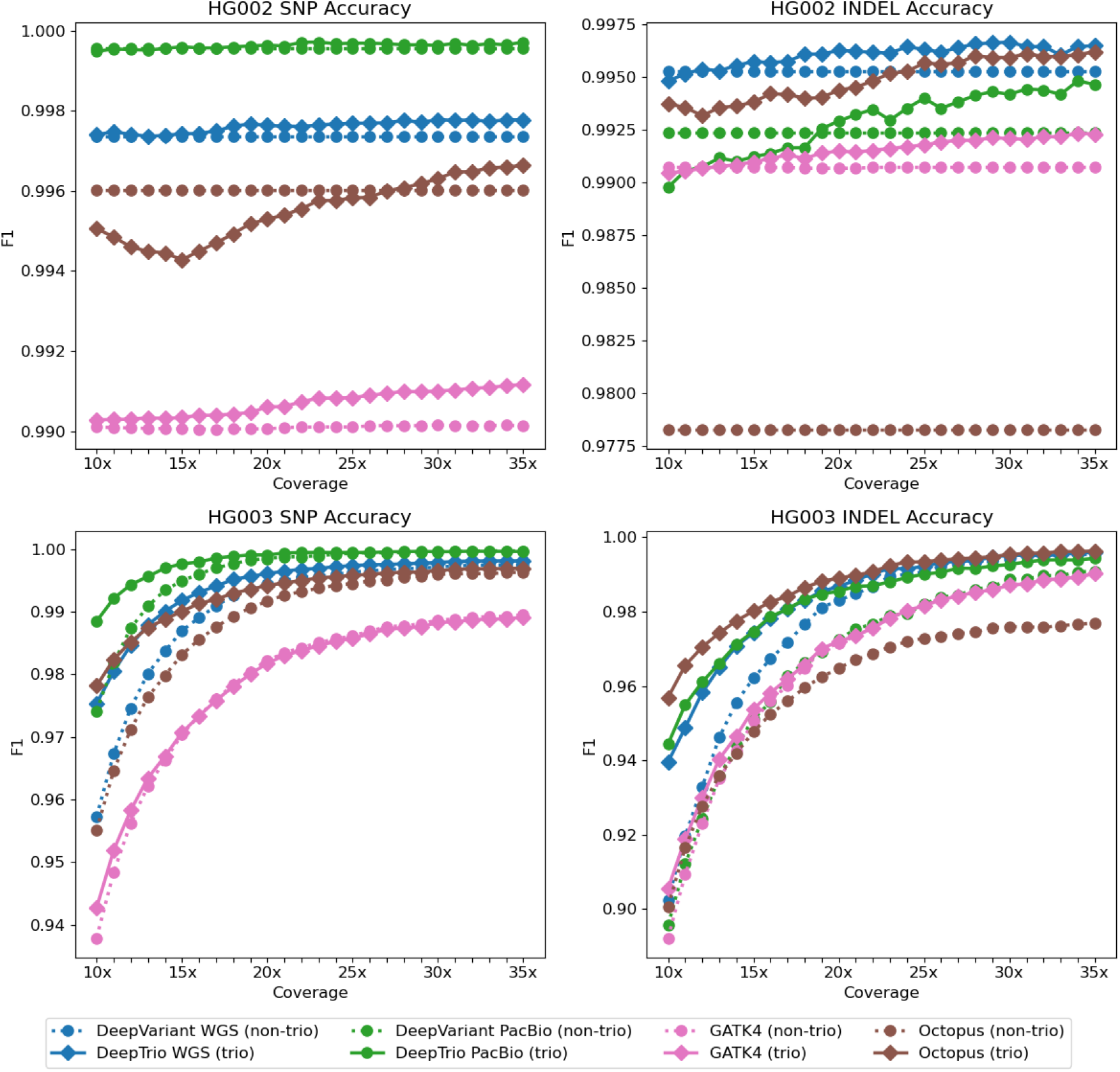
Variant calling accuracy of DeepTrio varying only parent coverage. Accuracy for DeepTrio and other pipelines over coverage titrations of the parent samples with the child sample at 35x. All samples use the same Illumina data, except for the green lines (DeepVariant PacBio and DeepTrio PacBio). Accuracy is determined by concordance with the Genome in a Bottle v4.2 truth set for the child (top) and parent (bottom). Trio methods are shown with a solid line and non-trio with a dotted line. F1 is determined from the total errors and correct calls for Indels added to SNPs for chromosome20. The F1 for dv-trio was much lower and was excluded.

**Figure 5.**
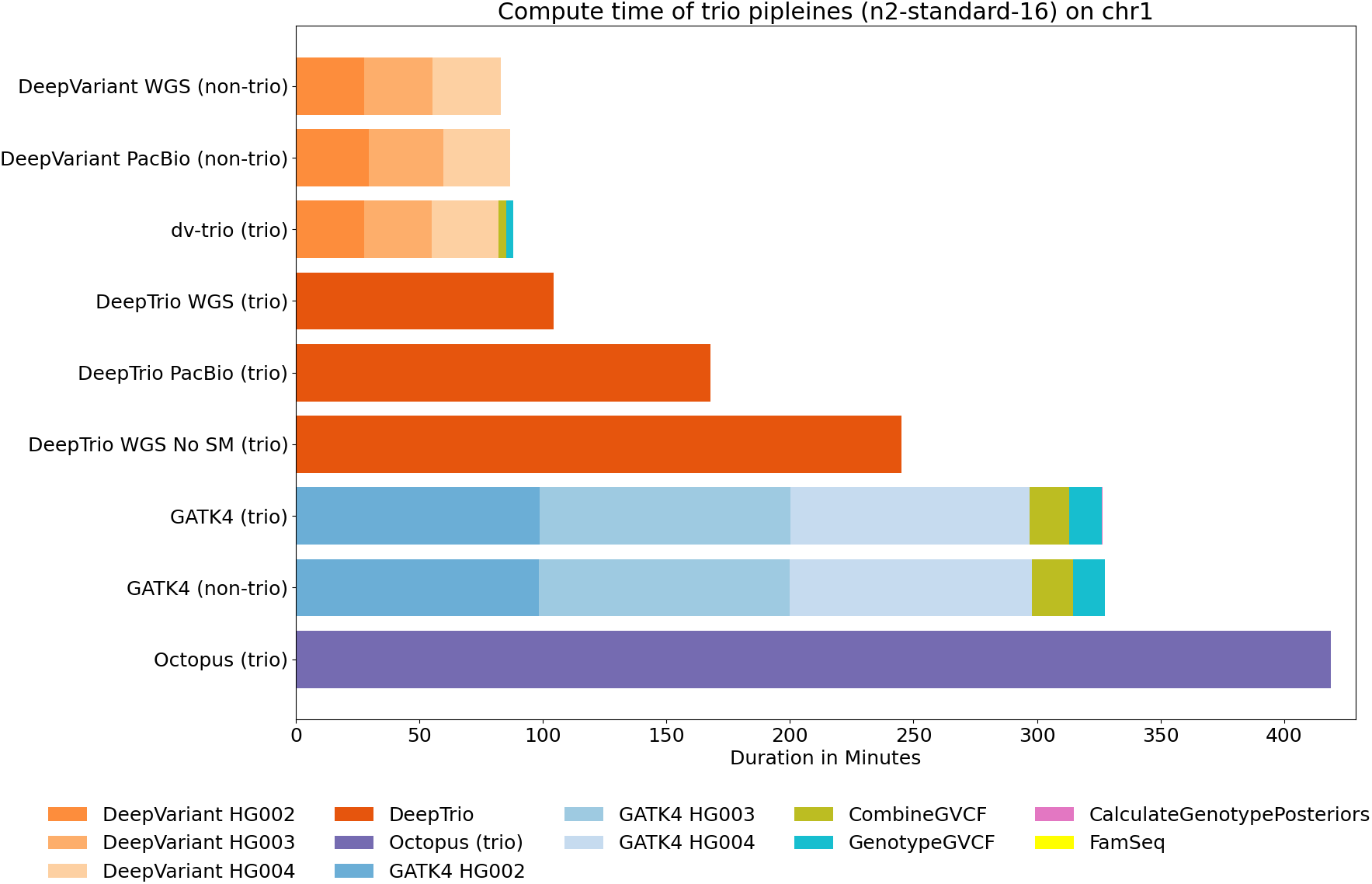
Time required to run DeepTrio and other pipelines. Compute time required for each stage of trio calling pipelines for a 35x Illumina WGS trio of HG002-HG003-HG004. Analysis is conducted on chromosome 1. The same machine type is used in each case, a 16-CPU thread instance *n1-standard-16*. Octopus (non-trio) was excluded due to its extensive runtime (3200 minutes total). DeepTrio without the use of the MLP (“No SM”) was included to illustrate its performance contributions.

**Figure 6.**
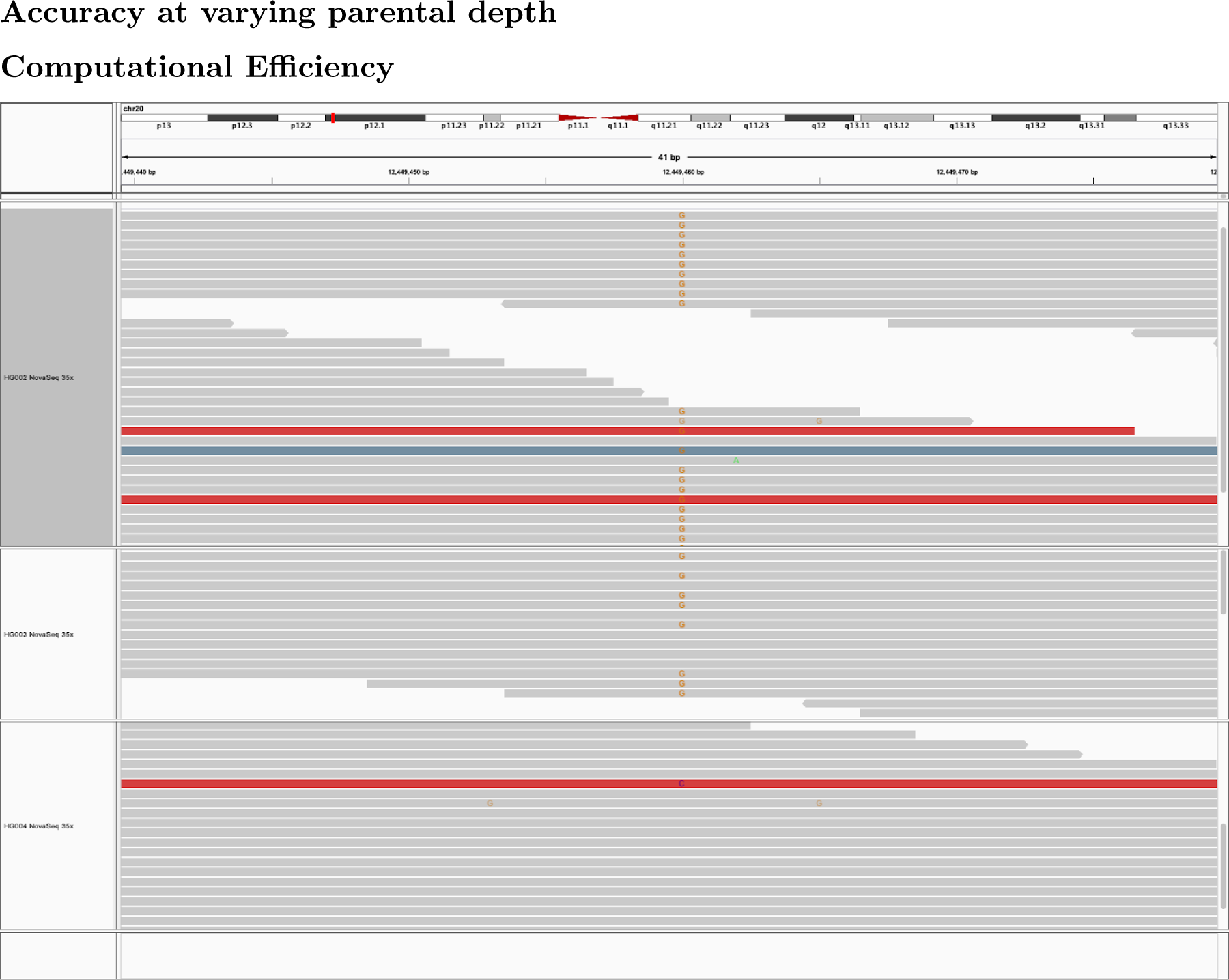
Example of a variant called correctly by DeepTrio but not DeepVariant IGV image of chr20:12449460 in 35x Illumina WGS PrecisionFDA v2 Truth Challenge samples for HG002-HG003-HG004. HG002 (child) is shown in the top row. This position is incorrectly genotyped as a homozygous variant in HG002 by DeepVariant and is correctly called a heterozygous variant in DeepTrio. This position is difficult to call because only a few reads are non-reference in HG002, and because some reads in this window are mapped discordantly.

**Figure 7.**
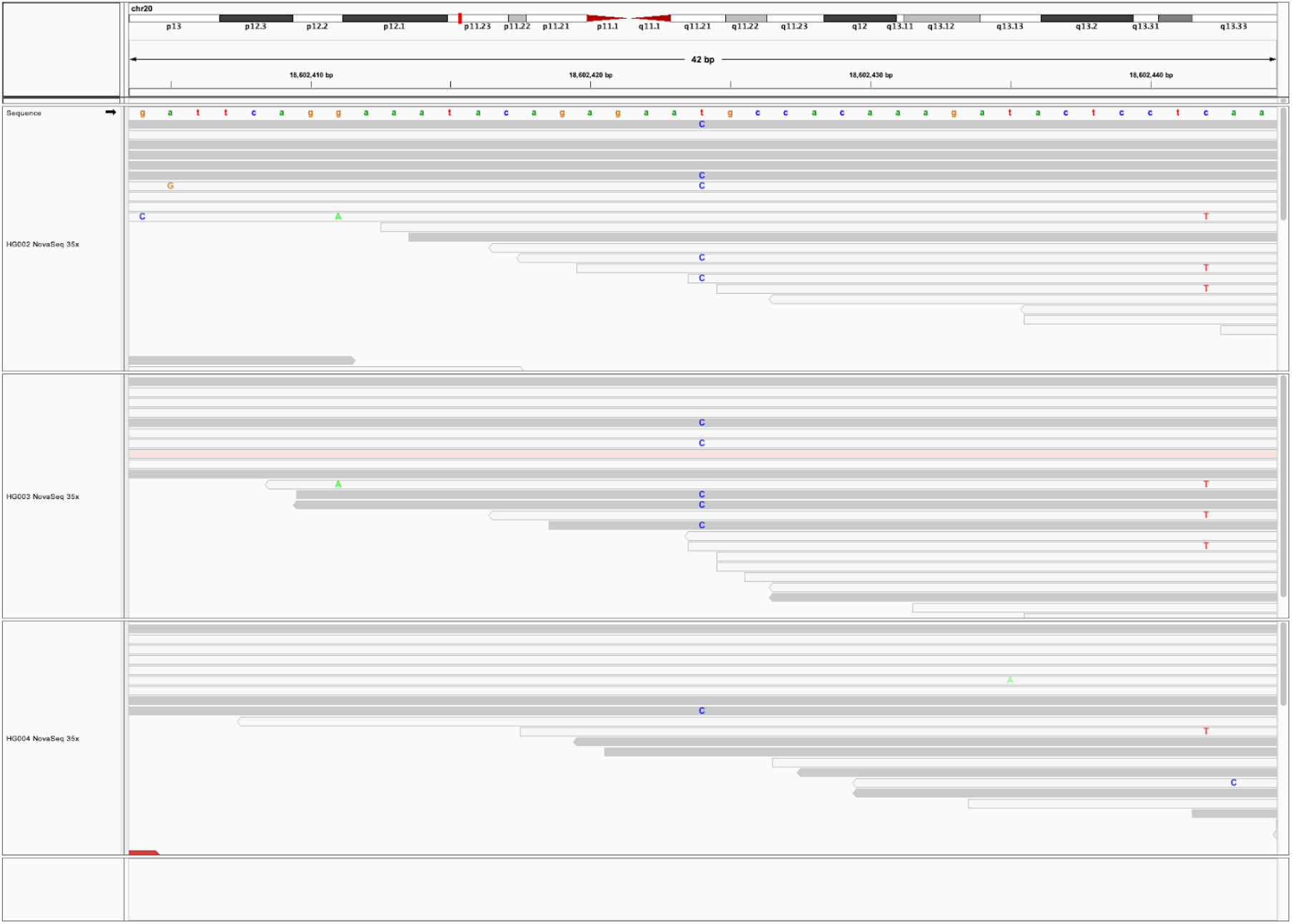
Example of a variant called correctly by DeepTrio but not DeepVariant. IGV image of chr20:18602424 in 35x Illumina WGS PrecisionFDA v2 Truth Challenge samples for HG002-HG003-HG004. HG002 (child) is shown in the top row. This position is called as a reference site in DeepVariant and is correctly called as a heterozygous variant in DeepTrio. The presence of this SNP in one of the parent samples strengthens credence to DeepTrio. IGV marks reads with MAPQ 0 as white instead of gray, indicating that this region is difficult to map with Illumina reads.

**Figure 8.**
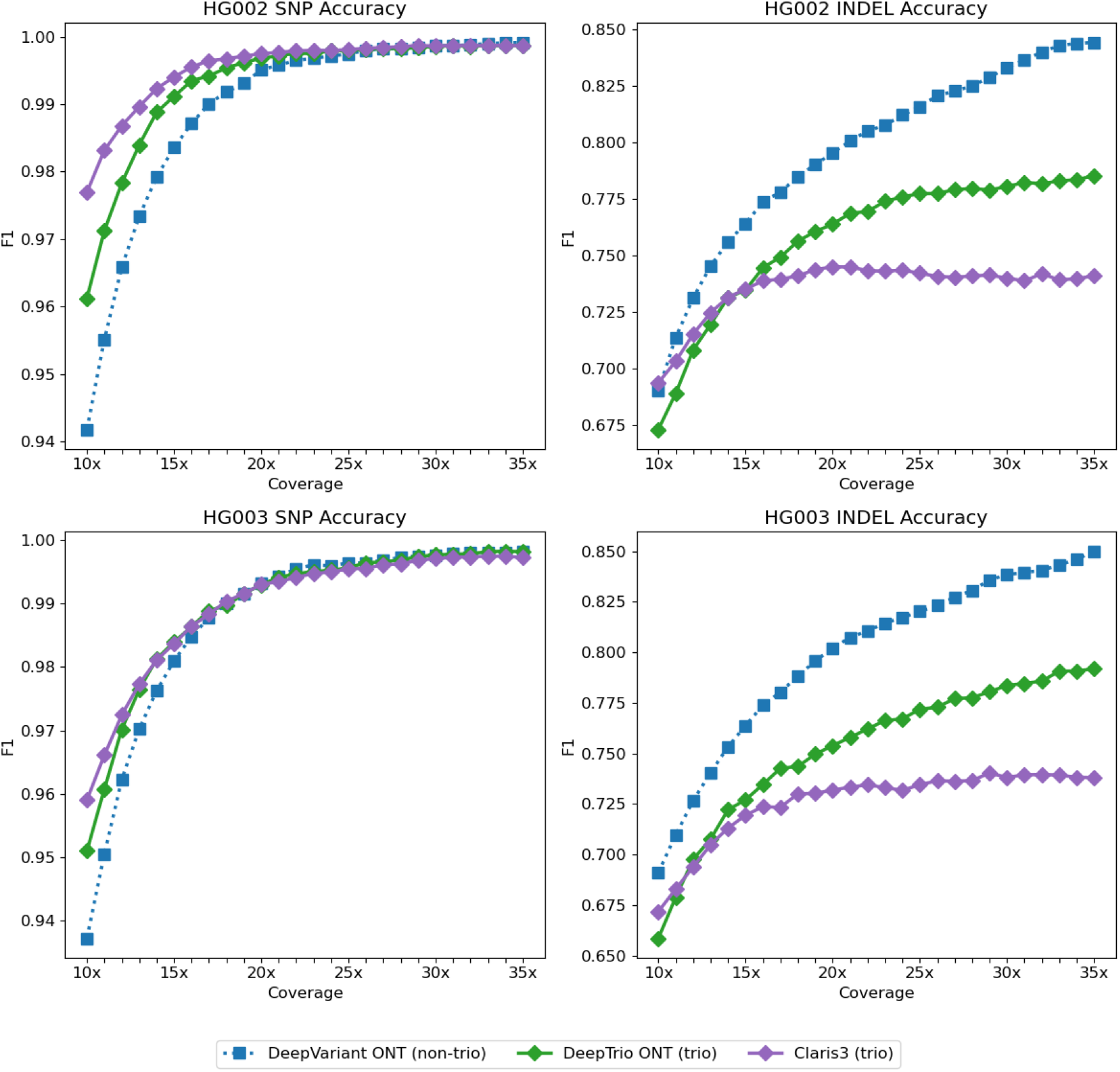
Variant calling accuracy for Oxford Nanopore long-reads. Accuracy for DeepTrio, DeepVariant and Clair3-trio over coverage titrations of the child and parent samples, determined by concordance with the Genome in a Bottle v4.2 truth set for the child (top) and parent (bottom), with the same coverage on all three samples. All samples use Oxford Nanopore Technology (ONT) long reads. F1 is determined from the total errors and correct calls for total Indels and SNPs for chromosome 20.

## Commands Used

**BWA**

~~~
bwa mem -t 16 references/grch38_bwa_index/genome.fa
 ${SAMPLE}.novaseq.pcr-free.${COVERAGE}x.R1.fastq.gz
 ${SAMPLE}.novaseq.pcr-free.${COVERAGE}x.R2.fastq.gz
 -R “@RG\\tID:${SAMPLE}\\tPL:ILLUMINA\\tPU:NONE\\tLB:${SAMPLE}\\tSM:${SAMPLE}”
| samtools sort -O BAM -o${SAMPLE}.novaseq.pcr-free.${COVERAGE}x.grch38.bam
~~~

**MarkDuplicates**

~~~
java -jar gatk-package-4.1.2.0-local.jar MarkDuplicates
 -I ${SAMPLE}.novaseq.pcr-free.${COVERAGE}x.grch38.bam
 -O ${SAMPLE}.novaseq.pcr-free.${COVERAGE}x.dedup.grch38.bam
 -M ${SAMPLE}.novaseq.pcr-free.${COVERAGE}x.dedup.grch38.metrics
~~~

**GATK HaplotypeCaller**

~~~
time sudo docker run \
 -v “${HOME}/input”:”/input” \
 -v “${HOME}/output”:”/output” \
 broadinstitute/gatk:${GATK_VERSION} gatk HaplotypeCaller \
 -I /input/$(basename “${READS_CHILD}”) \
 -R /input/$(basename “${REF}”) \
 -O /output/${SAMPLE_NAME_CHILD}.gatk.g.vcf.gz \
 ${REGIONS_FLAG} \
 --emit-ref-confidence GVCF
~~~

**GATK CombineGVCF**

~~~
time sudo docker run \
 -v “${HOME}/input”:”/input” \
 -v “${HOME}/output”:”/output” \ broadinstitute/gatk:${GATK_VERSION} gatk CombineGVCFs \
 -R /input/$(basename “${REF}”) \
 -V /output/${SAMPLE_NAME_CHILD}.gatk.g.vcf.gz \
 -V /output/${SAMPLE_NAME_PARENT1}.gatk.g.vcf.gz \
 -V /output/${SAMPLE_NAME_PARENT2}.gatk.g.vcf.gz \
 -O /output/${COMBINED_GVCF_NAME}
~~~

**GATK GenotypeGVCF**

~~~
time sudo docker run \
 -v “${HOME}/input”:”/input” \
 -v “${HOME}/output”:”/output” \ broadinstitute/gatk:${GATK_VERSION} gatk GenotypeGVCFs \
 -R /input/$(basename “${REF}”) \
 -V /output/${COMBINED_GVCF_NAME} \
-O /output/${GENOTYPED_VCF_NAME} \
--standard-min-confidence-threshold-for-calling 1
~~~

**GATK CalculateGenotypePosteriors**

~~~
time sudo docker run \
 -v “${HOME}/input”:”/input” \
 -v “${HOME}/output”:”/output” \
 broadinstitute/gatk:${GATK_VERSION} gatk CalculateGenotypePosteriors \
 -V /output/${GENOTYPED_VCF_NAME} \
 -ped /input/trio.ped \
 -O /output/${OUTPUT_VFC_NAME} \
 --skip-population-priors
~~~

**DeepVariant**

~~~
sudo docker pull google/deepvariant:${DEEPVARIANT_BIN_VERSION}
time sudo docker run \
 -v “${HOME}/input”:”/input” \
 -v “${HOME}/output”:”/output” \
 google/deepvariant:${DEEPVARIANT_BIN_VERSION} \
 /opt/deepvariant/bin/run_deepvariant \
 --model_type ${model_type} \
 --ref /input/$(basename “${REF}”) \
 --reads /input/$(basename “${reads}”) \
 --sample_name “${sample_name}” \
 --output_vcf /output/${sample_name}.vcf.gz \
 --output_gvcf /output/${sample_name}.g.vcf.gz \
 ${REGIONS_FLAG} \
 --num_shards “${NUM_THREADS}”
~~~

**Glnexus**

~~~
sudo docker run \
 -v “${HOME}/output”:”/output” \
 quay.io/mlin/glnexus:v1.2.7 /usr/local/bin/glnexus_cli \
 --config “DeepVariant_unfiltered” \
 “/output/${SAMPLE_NAME_CHILD}.g.vcf.gz” \
 “/output/${SAMPLE_NAME_PARENT1}.g.vcf.gz” \
 “/output/${SAMPLE_NAME_PARENT2}.g.vcf.gz” \
| bcftools view -O z -o “${HOME}/output/${TRIO_GVCF_NAME}”
~~~

**DeepTrio**

~~~
sudo docker pull google/deepvariant:deeptrio-${DEEPTRIO_BIN_VERSION}
time sudo docker run \
 -v “${HOME}/input”:”/input” \
 -v “${HOME}/output”:”/output” \
 google/deepvariant:deeptrio-${DEEPTRIO_BIN_VERSION} \
 /opt/deepvariant/bin/deeptrio/run_deeptrio \
 --model_type ${model_type} \
 --ref /input/$(basename “${REF}”) \
 --reads_child /input/$(basename “${READS_CHILD}”) \
 --reads_parent1 /input/$(basename “${READS_PARENT1}”) \
 --reads_parent2 /input/$(basename “${READS_PARENT2}”) \
 --output_vcf_child /output/${SAMPLE_NAME_CHILD}.vcf.gz \
 --output_vcf_parent1 /output/${SAMPLE_NAME_PARENT1}.vcf.gz \
 --output_vcf_parent2 /output/${SAMPLE_NAME_PARENT2}.vcf.gz \
 --sample_name_child “${SAMPLE_NAME_CHILD}” \
 --sample_name_parent1 “${SAMPLE_NAME_PARENT1}” \
 --sample_name_parent2 “${SAMPLE_NAME_PARENT2}” \
 --num_shards “${NUM_THREADS}” \
 $REGIONS_FLAG \
 --output_gvcf_child /output/${SAMPLE_NAME_CHILD}.g.vcf.gz \
 --output_gvcf_parent1 /output/${SAMPLE_NAME_PARENT1}.g.vcf.gz \
 --output_gvcf_parent2 /output/${SAMPLE_NAME_PARENT2}.g.vcf.gz
~~~

**dv-trio Famseq**

~~~
FamSeq vcf \
-vcfFile “${OUTPUT_DIR}/${GENOTYPED_VCF_NAME}.preprocessed.vcf” \
-pedFile “${PED_FILE}” \
-output “${OUTPUT_DIR}/${FAMSEQ_VCF_NAME}”
~~~

**Octopus (non-trio)**

~~~
time sudo docker run \
 -v “${HOME}/input”:”/input” \
 -v “${HOME}/output”:”/output” \
dancooke/octopus \
 -R input/${ref_name} \
 -I input/$(basename “${READS_CHILD}”) \
 ${REGIONS_FLAG} \
 --sequence-error-model PCR-FREE.NOVASEQ \
 --forest opt/octopus/resources/forests/germline.v0.7.4.forest \
 -o output/${SAMPLE_NAME_CHILD}.g.vcf.gz \
 --refcall BLOCKED \
 --threads $NUM_THREADS
~~~

**Octopus (trio)**

~~~
time sudo docker run \
 -v “${HOME}/input”:”/input” \
 -v “${HOME}/output”:”/output” \
dancooke/octopus \
 -R input/${ref_name} \
 -I input/$(basename “${READS_CHILD}”) input/$(basename “${READS_PARENT1}”) input/$(bas
 -M ${SAMPLE_NAME_PARENT1} \
 -F ${SAMPLE_NAME_PARENT2} \
 ${REGIONS_FLAG} \
 --sequence-error-model PCR-FREE.NOVASEQ \
 --forest opt/octopus/resources/forests/germline.v0.7.4.forest \
 -o output/${SAMPLE_NAME_CHILD}-${SAMPLE_NAME_PARENT1}-${SAMPLE_NAME_PARENT2}.g.vcf.gz
 --refcall BLOCKED \
 --threads $NUM_THREADS
~~~

**Clair3-trio**

~~~
time docker run \
 -v “${INPUT_DIR}”:”/input” \
 -v “${OUTPUT_DIR}”:”/output” \
hkubal/clair3-trio:${CLAIR3_TRIO_VERSION} \
 /opt/bin/run_clair3_trio.sh \
 --ref_fn=/input/${ref_name} \
 --bam_fn_c=/input/$(basename “${READS_CHILD}”) \
 --bam_fn_p1=/input/$(basename “${READS_PARENT1}”) \
 --bam_fn_p2=/input/$(basename “${READS_PARENT2}”) \
 --sample_name_c=“${SAMPLE_NAME_CHILD}” \
 --sample_name_p1=“${SAMPLE_NAME_PARENT1}” \
 --sample_name_p2=“${SAMPLE_NAME_PARENT2}” \
 --threads=${NUM_THREADS} \
 --model_path_clair3=“/opt/models/clair3_models/${MODEL_C3}” \
 --model_path_clair3_trio=“/opt/models/clair3_trio_models/${MODEL_C3T}” \
 --gvcf \
 --output=/output
~~~

**Accuracy Comparison with hap.py**

~~~
time sudo docker run \
 -v “${HOME}/input”:”/input” \
 -v “${HOME}/output”:”/output” \
jmcdani20/hap.py:v0.3.12 /opt/hap.py/bin/hap.py \
 “/input/$(basename “${gcs_truth_vcf}”)” \
 “/output/$(basename “${query_vcf_name}”)” \
 -r “/input/${ref_name}” \
 -f “/input/$(basename “${gcs_truth_bed}”)” \
 -o “/output/${sample}.happy” \
 -l “${CHROMOSOME}” \
 --pass-only \
 --no-roc \
 --no-json \
 --engine=vcfeval \
 --engine-vcfeval-template=/input/${ref_sdf_file} \
 --threads=${NUM_THREADS}
The truth VCF and BED files for Hap.py comparison are the v4.2.1 Truth Set from Genome in a Bottle: ftp://ftp-trace.ncbi.nlm.nih.gov/giab/ftp/release/AshkenazimTrio
~~~

